# Noninvasive optical monitoring of cerebral hemodynamics in a preclinical model of neonatal intraventricular hemorrhage

**DOI:** 10.1101/2024.10.16.618768

**Authors:** Jyoti V. Jethe, YuBing Y. Shen, Edmund F. LaGamma, Govindaiah Vinukonda, Jonathan A. N. Fisher

**Affiliations:** Department of Physiology, New York Medical College; Department Pediatrics, Division of Newborn Medicine, New York Medical College; Department of Biochemistry and Molecular Biology, New York Medical College; Department of Cell Biology and Anatomy, New York Medical College

**Keywords:** intraventricular hemorrhage, cerebral blood flow, hemodynamics, diffuse correlation spectroscopy, microvascular flow, cranial ultrasound, germinal matrix, pediatrics, autoregulation

## Abstract

Intraventricular hemorrhage (IVH) is a common complication in premature infants and is associated with white matter injury and long-term neurodevelopmental disabilities. Standard diagnostic tools such as cranial ultrasound and MRI are widely used in both preclinical drug development and clinical practice to detect IVH. However, these methods only provide endpoint assessments of blood accumulation and lack real-time information about dynamic changes in ventricular blood flow. This limitation could potentially result in missed opportunities to advance drug candidates that may have protective effects against IVH. In this pilot study, we aimed to develop a noninvasive optical approach using diffuse correlation spectroscopy (DCS) to monitor real-time hemodynamic changes associated with hemorrhagic and sub-hemorrhagic events in a preclinical rabbit model of IVH. DCS measurements were conducted during the experimental induction of IVH, and results were compared with ultrasound and histological analysis to validate findings. Significant changes in hemodynamics were detected in all animals subjected to IVH-inducing procedures, including those that did not show clear positive results on ultrasound. The study revealed progressively elevated coefficients of variation in blood flow, particularly driven by increased oscillations within the 0.05-0.1 Hz frequency band. These hemodynamic changes were more pronounced in animals that developed IVH, as confirmed by ultrasound. Our findings suggest that real-time optical monitoring with DCS can provide critical insights into pathological blood flow changes, offering a more sensitive and informative tool for evaluating potential therapeutics in the context of IVH.

## Introduction

There are roughly 15 million premature births worldwide every year, accounting for nearly 11% of all births [1]. Among the large number of other adverse health conditions, premature infants are susceptible to intraventricular hemorrhage (IVH) which is characterized by initial bleeding in the ganglionic eminence and subsequent accumulation of blood in the lateral ventricles. Reports indicate that roughly 45% of infants born before 32 weeks of gestation experience this condition [2]. Currently, no definitive treatment is available. This burden is particularly acute in low-income countries because of limited access to advanced neonatal care [3]. Premature infants are highly susceptible to the condition because a host of developmental vulnerabilities in the germinal matrix that collectively render the developing microvasculature mechanically unstable and prone to rupture. The mechanical vulnerability largely reflects developmental idiosyncrasies of the blood brain barrier, such as a paucity of pericytes, low levels of fibronectin in the basal lamina, and reduced expression of glial fibrillary acidic protein (GFAP) in astrocyte end feet [4].

While most cases of IVH in premature infants are diagnosed within 7 days after birth, the greatest risk for bleeding occurs within the first 48 hours of life [5]. If bleeding can be identified at an early stage, using preventive measures/timely intervention could help reduce the instability in germinal matrix vasculature. Early detection is crucial for not just minimizing further bleeding but also prevent the long-term impact on neurological development and support the infant’s physiological stability during this critical period. Currently, clinical methods for detecting IVH include cranial ultrasound and magnetic resonance imaging (MRI) [6]. Compared with other imaging modalities, ultrasound is by far the most used in clinical practice providing high-resolution images without ionizing radiation [6]. Its advantages include accessibility, portability, safety, and cost-effectiveness [7]. Limitations of the technique, however, include operator-dependent variability in imaging quality and findings. Subtle irregularities in small fontanelles, for example, may be missed by a novice technician. Additionally, ultrasound generally provides endpoint diagnostics rather than real-time monitoring. For functional measurements, Doppler ultrasonography can be used to detect blood flow [8]. However, the measurements are constrained to large vessels rather than microvasculature, which is the predominant source of rupture in IVH. While advanced imaging approaches such as arterial spin labeling (ASL) MRI could potentially detect such flow changes, the technique is certainly not realistic for constant monitoring of premature infants [9].

Diffuse optical technologies such as diffuse optical tomography (DOT) and near-infrared spectroscopy (NIRS) have found success in clinical adoption for real-time neuromonitoring of neonatal cerebral hemodynamics [10]. Pulse-DOT, for instance, has been used to measure cerebral hemodynamics in preterm infants including those who develop IVH [11]. Diffuse correlation spectroscopy (DCS) is another emerging optical method that has demonstrated promise for monitoring tissue hemodynamics in real-time [12]. DCS has been used effectively for non-invasive monitoring of cerebral hemodynamics and autoregulation in neurocritical care [13], and has also been applied to measurement of blood flow in infants with IVH [14]. DCS utilizes temporal fluctuations of light scattered by moving red blood cells to obtain a relative blood flow index, which units of cm^2^/s [15].

In this pilot study, we utilized a premature rabbit infant model to explore blood flow in the early phases of induced IVH. The glycerol induced IVH model of the rabbit pup brain closely mimics the developmental stages of human brains and reflects many of the clinical features of IVH such as hypomyelination, gliosis, pro-inflammatory cytokines, neurodegeneration, and cognitive delays seen in premature human neonates [16]. Beyond demonstrating sensitivity to IVH using this controlled model for the condition, we were able to detect sub-hemorrhagic hemodynamic changes that would otherwise go undetected. Our approach suggests that DCS could offer potentially sensitive biomarkers for preclinical exploration of new therapies, which is otherwise largely guided by terminal endpoints.

## Methods

### Surgical Procedures

All animal experiments were performed in accordance with the guidelines of the institutional Animal Care and Use Committee of New York Medical College. Timed pregnant New Zealand white rabbits were procured from Charles River Laboratories Inc (Wilmington, MA), whose pregnancies were carefully monitored. Premature pups were delivered via cesarean section at embryonic day 29 gestational age (rabbit full-term pregnancy is 32 days). Newborn premature pups, at 3–4 hours of postnatal age, were secured in a custom padded capsule (Fig. 1B) and treated with an intraperitoneal injection of 50% glycerol (6.5 g/kg, in physiological saline) which has been found to induce IVH in approximately 70% of treated animals [16]. Following approved ethical protocols, premature rabbit infants were not anesthetized. To determine the presence and severity of IVH, cranial ultrasound was performed at 24 hours postnatal age (Acuson Sequoia C256, Siemens Corp., Washington, DC).

**Figure 1:**
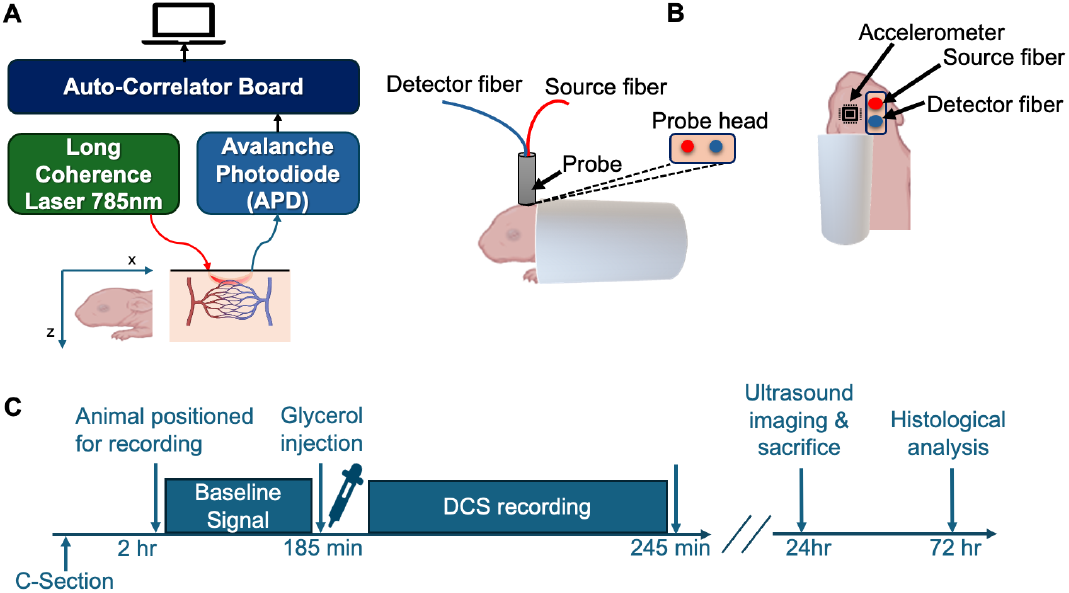
Experimental apparatus and protocol. **A** depicts a block diagram of the main components of the DCS system used in experiments. **B** illustrates the positioning of a premature rabbit infant and placement of the optical probe. Source-detector separation was 5 mm. **C** shows the experimental protocol and tissue processing timeline. Following acquisition of baseline flow signals for up to 25 min, glycerol (50%) was injected intraperitoneally. Optical signals were then continued recorded for another 60 min. Head ultrasound was performed 24 hours after birth, followed directly by sacrifice and perfusion for subsequent histological analysis.

### DCS System and Data Processing

The details of our DCS apparatus, depicted in Fig. 1A, have been described previously [17]. Briefly, 785-nm laser light from a long coherence length continuous-wave NIR laser (CrystaLaser, NV) was delivered to the animal’s scalp via multimode optical fiber (200-μm core diameter, Fiberoptic Systems Inc, CA). Reemitted light was relayed from the scalp to a photon counting avalanche photodiode (APD) (SPCM-AQRH-12-FC, Excelitas, Quebec, Canada) via a single-mode fiber (5-μm core diameter, Fiberoptic Systems Inc, California). The output of the APD was directed to an autocorrelator signal processing board (Correlator.com), which computes an intensity autocorrelation function that is continuously sent via universal serial bus (USB) to a laptop PC (Dell Precision 7510) running LabVIEW software (National Instruments, Austin, TX) for further data analysis. Using custom code in MATLAB (ver. R2022b, Mathworks, Inc., Waltham, MA), a tissue cerebral blood flow index (CBFi) is obtained by fitting the measured intensity correlation function to a correlation diffusion equation [12] using an absorption coefficient μ_a_ = 0.1 cm^−1^ and reduced scattering coefficient of μ_s_′ = 8.0 cm^−1^. Source and detection fibers were integrated into a custom probe that was 3D printed using thermoplastic polyurethane (TPU) material (Shapeways, Inc.). The source-detector fiber separation was 5mm. To monitor mechanical response and potential motion artifacts, the probe also integrated an accelerometer component (AST1001-BMA250, TinyCircuits) whose output was relayed to the same data acquisition laptop via USB. Individual CBFi measurements were acquired at 1 Hz and smoothed with 8s moving mean filter. The accelerometer sensor acquires data in three orthogonal axes: x, y, and z. To ensure comparability between the axes, the raw data from each axis was first normalized making it easier to interpret and analyze. Following normalization, the accelerometer data’s 3-dimentional magnitude was calculated by taking the square root of the sum of the squares of the acceleration values, i.e.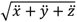, where 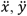, and 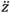 are the normalized acceleration measurements in each direction. Blood flow fluctuations were quantified in terms of the coefficient of variation (CV), defined as *σ*⁄*μ*, where *σ* is the standard deviation and *μ* is the mean. Motion analysis was quantified based on the CV of the 3-dimensional magnitude of accelerometry data (CV_*accel*_).

### Optical Phantom Experiments

To verify that optically-derived CBFi values and mechanical measurements were independent, we performed optical “phantom” experiments where phenomenological “flow” was controlled using optically detected Brownian motion in aqueous intralipid emulsions of varying viscosity [18]. A 1-gallon black polypropylene bucket was filled with a solution of water, glycerol, and intralipid. In this setup, a probe was inserted into the bucket, with an accelerometer attached to the probe allowing simultaneous acquisition of both flow signals and accelerometer data. High and low flow rates were simulated with glycerol concentrations of 2.26 M and 4.52 M, respectively.

### Experimental Protocol

After animals were positioned, the 3D-printed probe, which was stabilized on a mechanical isolation table with optomechanic posts (Thorlabs, NJ), was carefully lowered onto the animal’s scalp above the region of the forebrain. This scalp position corresponds to a region atop lateral ventricles on the mid-sagittal line and Bregma [16]. A total of six premature rabbit infants were selected as subjects for the experiment. The study involved 5 animals injected with glycerol and a sham animal that was positioned and measured identically but did not receive glycerol injection. As depicted in the timeline in Fig. 1D, following positioning in the measurement apparatus, hemodynamics were monitored for a period of 5 min, after which non-sham animals were injected with glycerol. Following the injection, optical measurements resumed for at least one hour. Animals were then returned to a neonatal incubator set at 27°C. A cranial ultrasound was obtained from animals 24 hours after the DCS recordings.

### Histological Analysis

To measure the ventricular cross-sectional area, coronal brain sections were prepared using hematoxylin-eosin (H&E) staining [16]. First, the rabbit pups were perfused with 1 × phosphate-buffered saline (PBS) buffer. Brains were dissected out and postfixed in 4% paraformaldehyde for 24h at 4 °C, after which they were dehydrated with 15% sucrose for 12h and then with 30% sucrose for 12h. Brains were then embedded in OCT compound (Tissue-Tek, Sakura Finetechnical Co., Tokyo, Japan) and sectioned into 18-μm thick coronal sections on a cryostat (CM1850, Leica Biosystems) [19].

The H&E staining procedure was performed in accordance with our previously published methods [16]. Briefly, sections were air-dried at room temperature (RT) for 20–30 min, followed by rinsing in PBS for 5 minutes (2 washes). Sections were fixed with acetic acid for 1 min, rinsed with water, and stained with hematoxylin for 1 minute. After an additional water rinse, sections were treated with ammonia water and rinsed again. They were then exposed to 95% ethanol for 15s, stained with eosin for 30s, and sequentially dehydrated in 95% ethanol (2 washes), followed by 100% ethanol, and cleared with three immersions in xylene. After staining the images were created using a microscope (Keyence, BZ-X810).

## Results

The experimental protocol and DCS apparatus for monitoring hemodynamics is shown in Fig. 1. Head ultrasound imaging performed 24 hours after birth validated IVH in 3 out of 5 injected animals based on increased echogenicity in the lateral ventricles (Fig. 2A-C). H&E staining of coronal brain sections from animals with IVH sacrificed directly following ultrasound imaging revealed extravasated erythrocytes within the ventricular lumen (Fig. 2D-F). In non-injected control animals, no increased echogenicity was observed in ultrasound (Fig. 2C) and the germinal matrix did not appear compromised (Fig. 2F).

**Figure 2:**
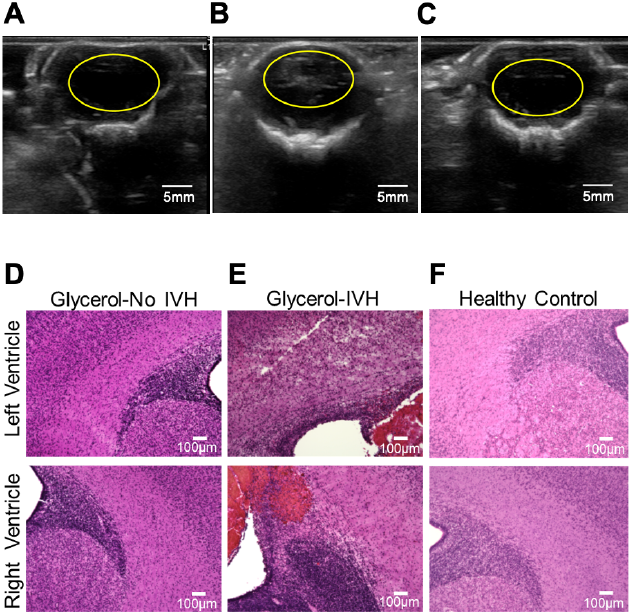
Validating IVH by means of ultrasound and tissue histology. **A-C** show representative ultrasound imaging results obtained 24 hours following experiments. The images correspond to representative IVH-, IVH+, and sham animals. In these cross-sectional images, the yellow ovals indicate the ventricular region. An example of increased echogenicity in the lateral ventricles can be seen within the yellow oval in **B. D-F** show representative images of 18-μm-thick H&E stained coronal brain slices from representative IVH-, IVH+, and sham animals, in sequence. Extravasated erythrocytes within the ventricular lumen can be seen in **E**.

Following glycerol injection, optically recorded hemodynamics waveforms were complex and varied among animals, as can be seen in the representative traces in Fig. 3A and B. Despite animal-to-animal variability in the raw blood flow signals, however, the coefficient of variation of cerebral blood flow (CV_*CBFi*_) increased following injection when summed over all animals. Fig. 3C depicts the CV_*CBFi*_ results of all animals, averaged in consecutive windows of 5 minutes, together with the results of linear regression that reveal a significant positive slope (*P* = 0.0004). Analyzing CV_*CBFi*_ separately on animals with and without IVH (IVH+ and IVH-, respectively), however, revealed that IVH+ animals demonstrated a greater and significant rate of increase over the course of 60 minutes following injection (*m* = 0.04, *P* = 0.0004) compared with IVH-animals (*m=* 0.01, *P* = 0.29) (Fig. 3D and E).

**Figure 3:**
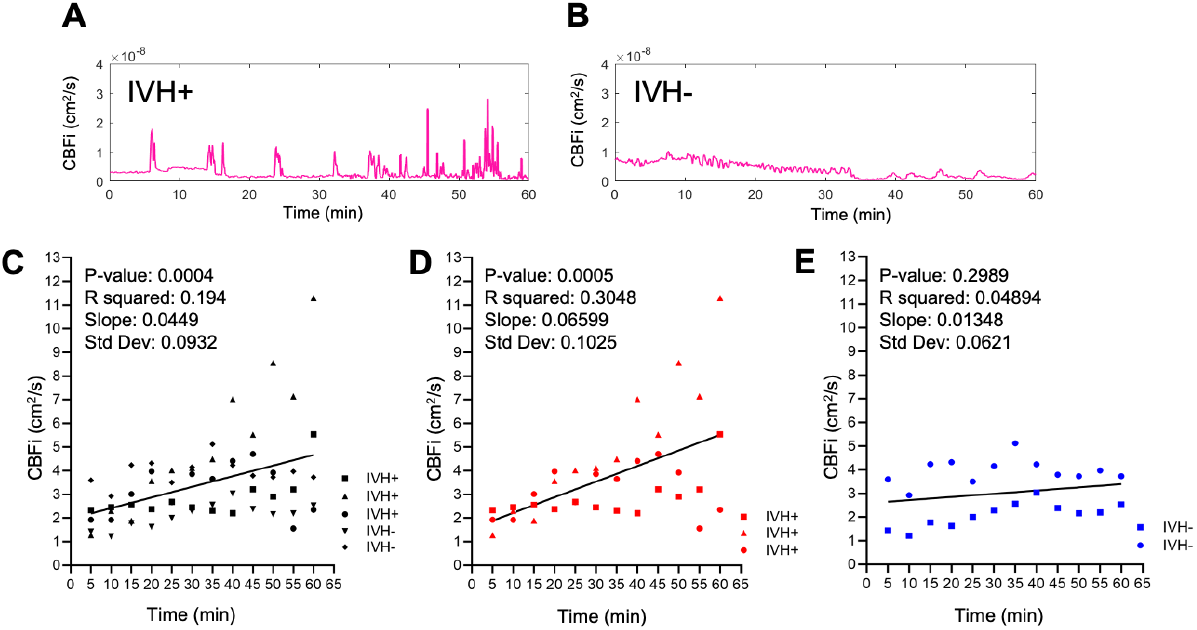
Optically-recorded cerebral blood flow index (CBFi) following glycerol injection. **A-B** depict flow recorded continuously for 60 min directly following IP injection of 50% glycerol. **C-E** depict coefficient of variation analysis and the superimposed lines show the results of linear regression. **C** shows the results of all non-sham animals (N = 5), and each data point represents CV_*CBFi*_ averaged over a 5-minute window. **D** shows the subset of data from IVH+ animals (N = 3), and **E** shows the data from only IVH-animals N = 2).

To explore whether apparent hemodynamic signals could have been a result of motion artifacts, we performed the same CV analysis on the 3-dimensional magnitude of accelerometry data (CV_*accel*_) and found no significant change over the course of the experiment for all animals (Fig. 4A). To verify that flow and accelerometry measurements were independent, we performed phantom measurements wherein flow was measured when the probe was immersed in highly scattering lipid emulsions of varying Newtonian viscosity. As shown in the scatterplot depicted in Fig. 4B, increasing viscosity reduced DCS-derived flow as expected because of the reduced Brownian motion of lipid scatterers. In contrast, the average motion as measured via accelerometry did not change (*P* = 0.83).

**Figure 4:**
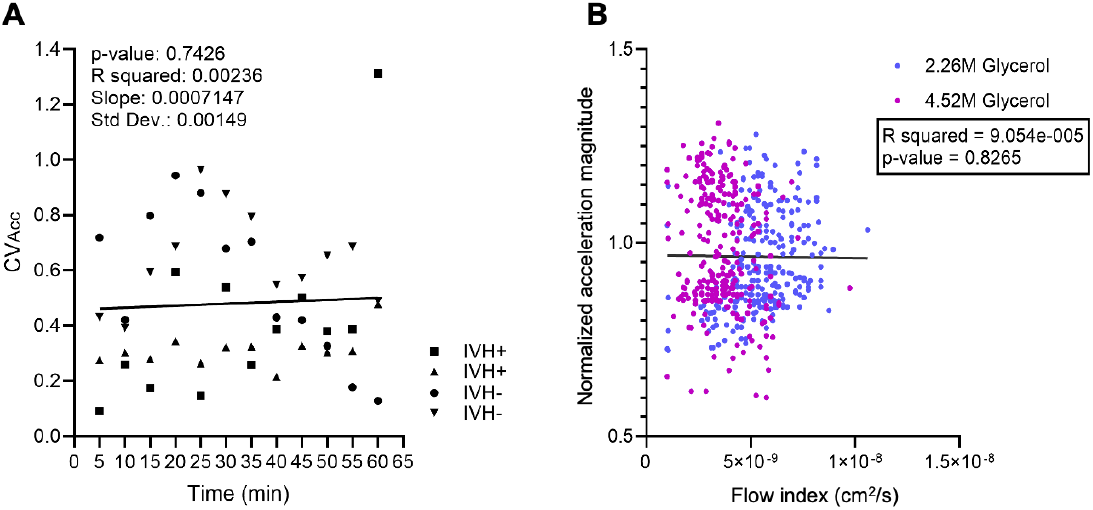
Physiological insult induces no significant motor or mechanical artifacts. **A** shows a plot of CV_*accel*_ for all experiments where accelerometry was possible (N = 4). Each data point represents CV_*accel*_ averaged over a 5 min window. Superimposed lines and statistics show the result of linear regression. **B** depicts the results of phantom measurements involving concurrent monitoring of 3D acceleration and DCS measurements of flow. The optical and mechanical sensors were bound and immersed in a highly-scattering intralipid solution at two viscosities. Each data point represents a single simultaneous measurement of 3D magnitude of acceleration paired with a CBFi at the same time point, and the integration time was 1.13s. All datapoints of one color are derived from a single experiment where data was acquired for 5 min.

## Discussion

Our main finding of heightened variability in CBFi following DCS likely reflects impaired autoregulation of cerebral blood flow owing to the significant physiological insult caused by the glycerol injection. In healthy animals, cerebral autoregulation ensures that cerebral blood flow is not heavily influenced by systemic blood pressure. Injecting glycerol interperitoneally results in dehydration and a high serum osmolarity; this been shown to induce selective rupture of vasculature in the germinal matrix [20]. The fluctuations that we observed, which were semi-periodic and largely within the frequency range of ∼0.05– 0.1 Hz, were consistent with dynamics that have been observed in a rabbit model of ischemic stroke [21].

Furthermore, increases in CV_*CBFi*_ variability began roughly 5 min post injection, which closely matches the onset of rapid increase in jugular venous pressure following IP injections of 50% glycerol in rats [22]. Given the vulnerability of microvasculature in the germinal matrix, we hypothesize that the fluctuations in CV_*CBFi*_ reflect flow dynamics in that vascular population.

While the CV_*CBFi*_ slope trend in IVH- animals did not achieve statistical significance within our sample size when analyzed separately from IVH+ animals, the difference in CV_*CBFi*_ trajectories may represent a reduced level of cerebral autoregulation impairment or else an early recovery from the physiological insult. The finding that glycerol injection did not elicit IVH in all animals is consistent with previous observations in this rabbit model [16]. Future experiments involving a larger sample size could help better resolve these trajectories and potentially distinguish the profiles of the two outcomes.

IVH studies in humans, albeit more directly clinically relevant than preclinical models, are challenging because the timeframe for developing IVH cannot be predicted and continual monitoring is not practical.

Prior work applying DCS to IVH in preterm infants, for example, utilized a case-control study design to retrospectively explore CBFi dynamics that were potentially predictive of IVH [23]. While elevated CV_*CBFi*_ was observed in infants who ultimately developed IVH, statistical significance of the correlation could not be achieved in those prior studies. This is likely because measurements were not obtained imminently before IVH, which spontaneous. Our preclinical model, however, facilitates a more direct path to developing quantitative real-time biomarkers because IVH is induced, and the timing has been demonstrated to be relatively predictable.

Overall, our proof-of-concept experiments have demonstrated that hemodynamics associated with the onset of IVH can be monitored in unanesthetized animals using an established preclinical model IVH. This could enhance the IVH drug development pipeline by providing a clinically relevant, real-time alternative to gold standard diagnostics such as ultrasound or histology which yield delayed, “binary” snapshots of the repercussions of IVH. Reliance on these existing approaches could therefore potentially lead to “no-go” decisions for otherwise promising drug design approaches that may offer potential protective effects.

